# Differential thermotolerance adaptation between species of *Coccidioides*

**DOI:** 10.1101/2020.08.12.247635

**Authors:** Heather L. Mead, Paris S. Hamm, Isaac N. Shaffer, Marcus de Melo Teixeira, Christopher S. Wendel, Nathan P. Wiederhold, George R. Thompson, Raquel Muñiz-Salazar, Laura Rosio Castañón-Olivares, Paul Keim, Carmel Plude, Joel Terriquez, John N. Galgiani, Marc J. Orbach, Bridget M. Barker

## Abstract

Coccidioidomycosis, or Valley fever, is caused by two species of dimorphic fungi. Based on molecular phylogenetic evidence, the genus *Coccidioides* contains two reciprocally monophyletic species: *C. immitis* and *C. posadasii.* However, phenotypic variation between species has not been deeply investigated. We therefore explored differences in growth rate under various conditions. A collection of 39 *C. posadasii* and 46 *C. immitis* isolates, representing the full geographical range of the two species, were screened for mycelial growth rate at 37°C and 28°C on solid media. The radial growth rate was measured over 16 days on yeast extract agar. A linear mixed effect model was used to compare the growth rate of *C. posadasii* and *C. immitis* at 37°C and 28°C respectively. *C. posadasii* grew significantly faster at 37°C, when compared to *C. immitis;* whereas both species had similar growth rates at 28°C. These results indicate thermotolerance differs between these two species. As the ecological niche has not been well-described for *Coccidioides* spp., and disease variability between species has not been shown, the evolutionary pressure underlying the adaptation is unclear. However, this research reveals the first significant phenotypic difference between the two species that directly applies to ecological and clinical research.

## 1. Introduction

Coccidioidomycosis, or Valley fever, is an environmentally acquired disease caused by inhalation of arthroconidia of dimorphic fungi belonging to the genus *Coccidioides*. In the environment, the fungi grow as filamentous mycelia, alternate cells of which autolyze and become fragile, leaving intact asexual arthroconidia that may disperse via wind or soil disruption. If inhaled by a susceptible host, an arthroconidium switches to a host-associated lifecycle and develops into a specialized infectious structure called a spherule. Subsequently, the host’s immune system either represses spherule replication or the host succumbs to the illness (1, 2). It is thought that symptomatic infection occurs in approximately 40% of human patients, who exhibit a broad spectrum of clinical symptoms, ranging from acute self-limited pneumonia, fibrocavitary chronic pulmonary infection, or hematogenous spread to extrapulmonary locations (i.e. disseminated infection) (3). By one estimate, there are 146,000 new symptomatic U.S. coccidioidal infections each year (4) although the reported cases are substantially lower (5).

Coccidioidomycosis is caused by two species, *C. immitis* and *C. posadasii.* Genetic analysis of multiple molecular markers has defined two monophyletic clades (6). Subsequent population genetic/genomic studies revealed that *C. immitis* is composed of at least two populations in the western U.S., and *C. posadasii* is composed of three populations widely dispersed across the American continents (7–10). Given the high number of autapomorphic mutations between *Coccidioides* species and among isolates within species, variation in phenotypes is predicted (11). However, minimal work characterizing phenotypic differences has been undertaken. A previous study demonstrated that *C. immitis in vitro* spherules grew in a synchronous pattern where *C. posadasii* isolates did not (12). Differences in pathogenesis and other disease-associated phenotypic characteristics among strains have been reported, although only one study had species information (13–18). The publication that defined the novel species *C. posadasii* also found species-specific variance in growth rate on media containing 0.136M NaCl, suggesting that *C. immitis* is more salt tolerant than *C. posadasii*, but due to overlap in the phenotype, and evaluation of only 10 isolates of each species, it was not statistically meaningful (6). These data supported observations published in the 1950s – 60s, which proposed that salinity of the soil may be a factor in determining the distribution of *C. immitis* in Californian soil (19­21). In contrast, a correlation of *C. posadasii* with saline soils was not observed in Arizona, where other associations were observed (22–26). Importantly, recent modeling analysis predicts the future expansion of *Coccidioides* species in response to climate dynamics (27). Therefore, a robust investigation of abiotic tolerances that may either limit or enhance distribution of *Coccidioides* is needed (1, 28, 29). Such vital information could provide clues regarding the ecological niche, geographical range limits, or host-specific adaptations of the two species of *Coccidioides*.

The division of *Coccidioides* into two species has been challenged by clinicians because of the lack of apparent difference in disease manifestation caused by the two pathogens, but recent work suggests that there might be differences in dissemination patterns between the species (1, 2, 30). Unfortunately, diagnosis and treatment of coccidioidomycosis does not require clinicians to identify to species. The current diagnostic methods; AccuProbe® (31), CocciDx (32), and CocciENV (33), do not distinguish between the two species. Molecular-based technologies exist to differentiate the two species, but these have not been adapted to clinical use (34, 35). However, genotyping the causative agent would allow correlation of clinical presentations and outcomes associated with species. Severe disease and death typically occurs in high risk group patients; however, seemingly healthy individuals can succumb as well, without a known host immunologic or pathogen genotypic explanation (36). Currently, the range of disease manifestations is suggested to be primarily due to host factors (37, 38). There are data supporting variation of virulence among individual isolates, but there is limited research on the subject (1, 13, 16, 17, 39). A reasonable hypothesis would acknowledge that both host and pathogen genetics play a role in disease outcome and should be further investigated (40–43). *Coccidioides,* like other primary fungal pathogens has evolved to withstand 37°C, mammalian body temperature which contributes to establishing host infection (44, 45).

This phenomenon, thermotolerance is an intrinsic characteristic of an organism that allows for tolerance of excessively high temperatures. Heat acclimation can shape natural populations for a wide range of microorganisms, and is a physiological adaptation to heat stress imposed by the colonization of new habitats, global climate change and encountering new hosts (46–54). This “preadaptation” is particularly important to pathogenic fungi that tolerate growth in high temperatures, which allows colonization of mammalian tissues (55, 56). For example, *Coccidioides* is adapted to grow at high temperatures in the environment (i.e. North and South American deserts), and is able to colonize a wide range of endothermic hosts throughout the Americas (57–61). C. immitis is endemic to the California Central Valley, whereas C. posadasii is widely distributed, but has highest prevalence in the Sonoran Desert. The annual mean temperature varies between the hotspot areas, with the California Central Valley having more mild temperatures compared to the Sonoran Desert, which led us to hypothesize that C. posadasii is more thermotolerant than C. immitis. Therefore, we investigated the growth rate of both species at 37°C representing host temperature and 28°C to support environmental growth conditions, so that we might elucidate species-specific phenotypic variation. Here we demonstrate thermotolerance dissimilarity of the two species by analyzing growth rates of 85 isolates at these two temperatures.

## 2. Materials and Methods

Strains and Media. 39 *C. posadasii* strains and 46 *C. immitis* strains used in this study are primarily human patient isolates archived by various institutions, as detailed in Table 1 (6, 8, 28, 62). These strains represent both the full geographic range of the two species, and the proposed geographically distinct sub-populations (6, 8). Strains were grown on 2xGYE media (2% glucose, 1% yeast extract, 1.5% agar w/v) to supply initial plugs to inoculate plates for growth analysis. Yeast Extract (YE) media (0.5% yeast extract, 1.5% agar w/v) was used for growth experiments. Flagstaff Medical Center isolates were collected under IRB No. 764034 through Northern Arizona Healthcare as part of the Northern Arizona University Biobank.

**Table 1.** Strain information

| ID | Species | Geographical Origin<sup>a</sup> | Source | Testing Institution |
| --- | --- | --- | --- | --- |
| CA22 | *C. immitis* | California | University of Texas Health Science Center (UTHSC) | NAU |
| 500 | *C. posadasii* | Soil, Tucson, AZ | University of Arizona (UA) | UA |
| IL1 | *C. posadasii* | Illinois | UTHSC | NAU |
| CA23 | *C. immitis* | California | UTHSC | NAU |
| HS-I-000718 | *C. posadasii* | Arizona | Flagstaff Medical Center (FMC) | NAU |
| GT164 | *C. posadasii* | Texas | University of California Davis (UCD) | NAU |
| GT163 | *C. immitis* | California | UCD | NAU |
| HS-I-000588 | *C. posadasii* | Arizona | FMC | NAU |
| CA28 | *C. immitis* | California | UTHSC | NAU |
| TX4 | *C. posadasii* | Texas | UTHSC | NAU |
| HS-I-000235 | *C. posadasii* | Arizona | FMC | NAU |
| TX1 | *C. posadasii* | Texas | UTHSC | NAU |
| HS-I-000778 | *C. posadasii* | Arizona | FMC | NAU |
| GT147 | *C. immitis* | California | UCD | NAU |
| HS-I-000234 | *C. posadasii* | Texas | FMC | NAU |
| CA30 | *C. immitis* | California | UTHSC | NAU |
| HS-I-000547 | *C. posadasii* | Arizona | FMC | NAU |
| HS-I-000233 | *C. posadasii* | Arizona | FMC | NAU |
| GT166 | *C. posadasii* | Texas | UCD | NAU |
| CA24 | *C. immitis* | California | UTHSC | NAU |
| CA29 | *C. immitis* | California | UTHSC | NAU |
| M211 | *C. posadasii* | Central Mexico | Unidad de Micologia,<br>UNAM | NAU |
| GT158 | *C. posadasii* | Arizona | UCD | NAU |
| CA15 | *C. immitis* | California | UTHSC | NAU |
| CA27 | *C. immitis* | California | UTHSC | NAU |
| TX3 | *C. posadasii* | Texas | UTHSC | NAU |
| CA20 | *C. immitis* | California | UTHSC | NAU |
| RS | *C. immitis* | California | Common Laboratory Strain | NAU |
| Silveira | *C. posadasii* | California | Common Laboratory Strain | NAU |
| RMSCC2378 | *C. posadasii* | Argentina | R. Negroni | UA |
| RMSCC2377 | *C. posadasii* | Argentina | R. Negroni | UA |
| RMSCC2379 | *C. posadasii* | Argentina | R. Negroni | UA |
| RMSCC3698 | *C. immitis* | Barstow, California | Naval Hospital | UA |
| RMSCC3490 | *C. posadasii* | Coahuila, Mexico | I. Gutierrez | UA |
| RMSCC3505 | *C. immitis* | Coahuila, Mexico | I. Gutierrez | UA |
| RMSCC3506 | *C. posadasii* | Coahuila, Mexico | I. Gutierrez | UA |
| RMSCC3472 | *C. posadasii* | Michoacán, Mexico | I. Gutierrez | UA |
| RMSCC3474 | *C. immitis* | Michoacán, Mexico | I. Gutierrez | UA |
| RMSCC3475 | *C. immitis* | Michoacán, Mexico | I. Gutierrez | UA |
| RMSCC3476 | *C. immitis* | Michoacán, Mexico | I. Gutierrez | UA |
| RMSCC3478 | *C. posadasii* | Michoacán, Mexico | I. Gutierrez | UA |
| RMSCC3479 | *C. immitis* | Michoacán, Mexico | I. Gutierrez | UA |
| RMSCC3377 | *C. immitis* | Monterey,<br>California | UCD | UA |
| RMSCC2343 | *C. posadasii* | Nuevo Leon,<br>Mexico | R. Diaz | UA |
| RMSCC2346 | *C. posadasii* | Nuevo Leon,<br>Mexico | R. Diaz | UA |
| RMSCC3738 | *C. posadasii* | Piaui, Brazil | B. Wanke | UA |
| RMSCC3740 | *C. posadasii* | Piaui, Brazil | B. Wanke | UA |
| RMSCC2127 | *C. posadasii* | Texas | UTHSC | UA |
| RMSCC2133 | *C. posadasii* | Texas | UTHSC | UA |
| RMSCC2234 | *C. posadasii* | Texas | UTHSC | UA |
| RMSCC2102 | *C. immitis* | San Diego,<br>California | University of California San<br>Diego (UCSD) Medical<br>Center | UA |
| RMSCC2394 | *C. immitis* | San Diego,<br>California | UCSD Medical Center | UA |
| RMSCC2395 | *C. immitis* | San Diego,<br>California | UCSD Medical Center | UA |
| RMSCC3693 | *C. immitis* | San Diego, California | Naval Hospital | UA |
| RMSCC3703 | *C. immitis* | San Diego, California | UCSD Medical Center | UA |
| RMSCC3705 | *C. immitis* | San Diego, California | UCSD Medical Center | UA |
| RMSCC3706 | *C. immitis* | San Diego, California | UCSD Medical Center | UA |
| RMSCC2006 | *C. immitis* | San Joaquin Valley | Kern County Public Health (KCPH) | UA |
| RMSCC2009 | *C. immitis* | San Joaquin Valley | KCPH | UA |
| RMSCC2010 | *C. immitis* | San Joaquin Valley | KCPH | UA and NAU |
| RMSCC2011 | *C. immitis* | San Joaquin Valley | KCPH | UA |
| RMSCC2012 | *C. immitis* | San Joaquin Valley | KCPH | UA |
| RMSCC2014 | *C. immitis* | San Joaquin Valley | KCPH | UA |
| RMSCC2015 | *C. immitis* | San Joaquin Valley | KCPH | UA |
| RMSCC2017 | *C. immitis* | San Joaquin Valley | KCPH | UA |
| RMSCC2268 | *C. immitis* | San Joaquin Valley | KCPH | UA |
| RMSCC2269 | *C. immitis* | San Joaquin Valley | KCPH | UA |
| RMSCC2271 | *C. immitis* | San Joaquin Valley | KCPH | UA |
| RMSCC2273 | *C. immitis* | San Joaquin Valley | KCPH | UA |
| RMSCC2274 | *C. immitis* | San Joaquin Valley | KCPH | UA |
| RMSCC2275 | *C. immitis* | San Joaquin Valley | KCPH | UA |
| RMSCC2276 | *C. immitis* | San Joaquin Valley | KCPH | UA |
| RMSCC2277 | *C. immitis* | San Joaquin Valley | KCPH | UA |
| RMSCC2278 | *C. immitis* | San Joaquin Valley | KCPH | UA |
| RMSCC2279 | *C. immitis* | San Joaquin Valley | KCPH | UA |
| RMSCC2280 | *C. immitis* | San Joaquin Valley | KCPH | UA |
| RMSCC2281 | *C. immitis* | San Joaquin Valley | KCPH | UA |
| RMSCC3480 | *C. posadasii* | Sonora, Mexico | I. Gutierrez | UA |
| RMSCC3487 | *C. posadasii* | Sonora, Mexico | I. Gutierrez | UA |
| RMSCC3488 | *C. posadasii* | Sonora, Mexico | I. Gutierrez | UA |
| RMSCC1040 | *C. posadasii* | Tucson, Arizona | UA | UA |
| RMSCC1043 | *C. posadasii* | Tucson, Arizona | UA | UA |
| RMSCC1044 | *C. posadasii* | Tucson, Arizona | UA | UA |
| RMSCC1045 | *C. posadasii* | Tucson, Arizona | UA | UA |
| RMSCC3796 | *C. posadasii* | Venezuela | G. San-Blas |  |

<sup>a</sup>Often patient diagnosis location

Growth Conditions and Measurements. Colonies were started by spreading approximately 10^6^ arthroconidia over the entire surface of a 2xGYE plate to create a lawn of mycelium to be transferred to initiate the thermotolerance experiment; this allowed measurement of colonial growth and not spore germination differences. After five days of growth at 25°C, 7mm diameter mycelial plugs were subcultured to the center of YE plates using a transfer tool (Transfertube® Disposable Harvesters, Spectrum® Laboratories). Three replicates of each strain were plated for each experiment. All plates (100mm x 15mm BD Falcon 1015) were sealed with gas permeable seals (labtape form TimeMed Labeling Systems, Inc or Key Scientific plate seals) for safety. Plates were placed in temperature-controlled incubators at either 28°C or 37°C in the dark under ambient humidity (30-50% RH) and CO_2_ (0.1%) conditions. Plate stacks were rotated from top to bottom and repositioned in the incubator with each measurement timepoint to reduce effects of environmental variation within the incubators. For measurement of radial growth, the diameter of each colony was measured in mm at 5, 7, 9, 12, 14, and 16 days post-subculture. The initial experiment proceeded at University of Arizona (UA) and subsequent testing with a new set of isolates occurred at Northern Arizona University (NAU). Details for strains tested at each institution are listed in Table 1 and all raw measurement data are available in File S1.

Statistical Analysis. To estimate the mean growth rate for each species over the two-week period a mixed effect linear model for each temperature was constructed using the lme4 package in R version 3.6.2 (63, 64). Initially, data sets were divided by institution and after concluding that species specific growth rate was not impacted by collection site the data sets were combined (Table S1). In the temperature specific models, the factors “day” and “species” were assumed to be fixed linear effects, and individual isolate response for each day was considered to be a normally distributed random effect as appropriate in a longitudinal study. Thus, the response variable of colony diameter was modeled with fixed effects and a random effect to determine if growth rates varied between strains at either 28°C and 37°C. Shapiro-Wilk test (p-value < 0.001) shows that residuals are not normally distributed. However, the large sample size and overall residual structure support that a linear model is the most appropriate for this data set. In addition, bootstrapping using the boot package in R (65, 66) was used to estimate 95% confidence intervals (CIs) for growth rates and other fixed effects (nsim=2,000). All bootstrap parameters were similar and support model estimates. A comparison between bootstrapped CIs and CIs constructed using the linear model can be found in S1 Table and S2 Table.

## 3. Results

To define variability of one phenotypic trait between two *Coccidioides* species, we examined the ability of *Coccidioides* spp. to grow in filamentous form at 37°C and 28°C on yeast extract (YE) agar. In this study, we surveyed 85 strains of *Coccidioides*, representing isolates from the entire geographical range of *Coccidioides,* for growth rate differences between species at 37°C and 28°C (Figure 1). Initial investigations occurred at the University of Arizona, and subsequent studies occurred at Northern Arizona University (Table 1).

**Fig 1.**
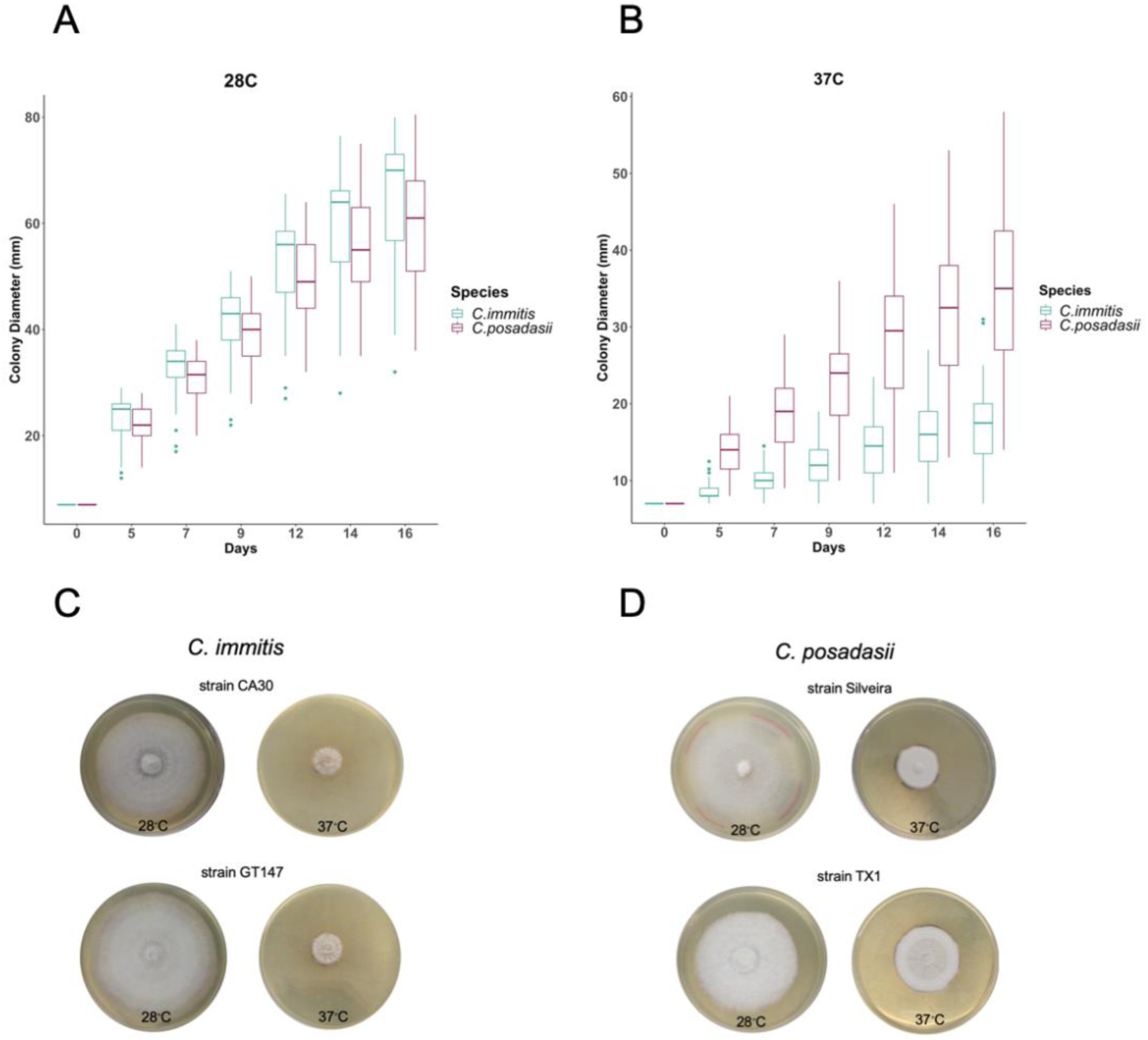
Temperature impacts growth ability of *C. immitis* isolates compared to *C. posadasii* on YE media. Seven mm diameter plugs were sub-cultured onto yeast extract plates and radial growth was documented over 16 days. (A) Radial growth measurements at 37°C for 46 *C. posadasii* and 39 *C. immitis* isolates in triplicate. (B) Radial growth measurements at 28°C for 46 *C. posadasii* and 39 *C. immitis* isolates in triplicate. (C) Representative samples of phenotypic variation observed between species on day 16.

We observed that mean growth rates varied slightly between institutions however overall species-specific temperature behavior remained. Therefore, data sets were combined (Figure S1, Table S1). Using a mixed effect linear model, we showed a significant species-specific difference for growth of the mycelial phase of the fungus based on temperature (Figure 2 and Table 2). Table 2 summarizes the estimated growth rate for each species, 95% confidence interval (CI), and p-value for each temperature specific model. Both species grew quicker at 28°C than 37°C. Although, *C. posadasii* had a larger mean diameter on all days tested (Table S3) the overall rate of increase was not statistically significant (p-value = 0.072, Table 2). This was in contrast to growth at 37°C. At this temperature, *C. posadasii* strains exhibited larger mean diameters, which reached double the diameter of *C. immitis* by day 16 (Table S3). At this temperature the overall growth rate of *C. posadasii* was 1mm/day faster than *C. immitis* (Figure 2 and Table 2). This difference was statistically significant (p-value < 0.001, Table 2). These findings were consistent for all days tested, and represent differential phenotypes for both species. Thus, our analysis indicates that high temperature is the important variable between species growth rate on solid media. This phenotypic difference supports the molecular phylogenetic species designation and may reflect adaptation of *C. immitis* to cooler environments, or possibly specific hosts.

**Table 2.** Temperature Specific Linear Model Slope Estimates for Radial Growth Rate at 28°C or 37°C.

| Species | Colony Diameter at 28°C |  |  | Colony Diameter at 37°C |  |  |
| --- | --- | --- | --- | --- | --- | --- |
|  | mm/day | 95% CI | *p*<sup>a</sup> | mm/day | 95% CI | *p*<sup>a</sup> |
| *C. immitis* x Day | 3.73 | 3.53 – 3.92 | 0.072 | 0.64 | 0.51 – 0.78 | <0.001 |
| *C. posadasii* x Day | 3.47 | 0.55 – 0.02 |  | 1.82 | 0.98 – 1.38 |  |
| N<sup>b</sup> | 85 |  |  | 85 |  |  |

<sup>a</sup> difference between estimated slope <sup>b</sup> number of isolates

Summary of temperature specific linear models, for 28°C and 37°C, respectively. Colony growth estimates for each species per day (slope), 95% confidence intervals (CI) and p values. At 28°C, *C. immitis* grows 3.73 mm/day which is 0.26 mm faster per day than *C. posadasii.* The difference in slope is not significant (p= 0.072) based on a=0.05. However, the p-value trends towards significance. At 37°C, *C. immitis* grows 0.64mm/day which is 1.18mm slower than *C. posadasii.* The difference in slope (CI, 0.98-1.38 mm/day) is statistically significant (p<0.001).

**Fig 2.**
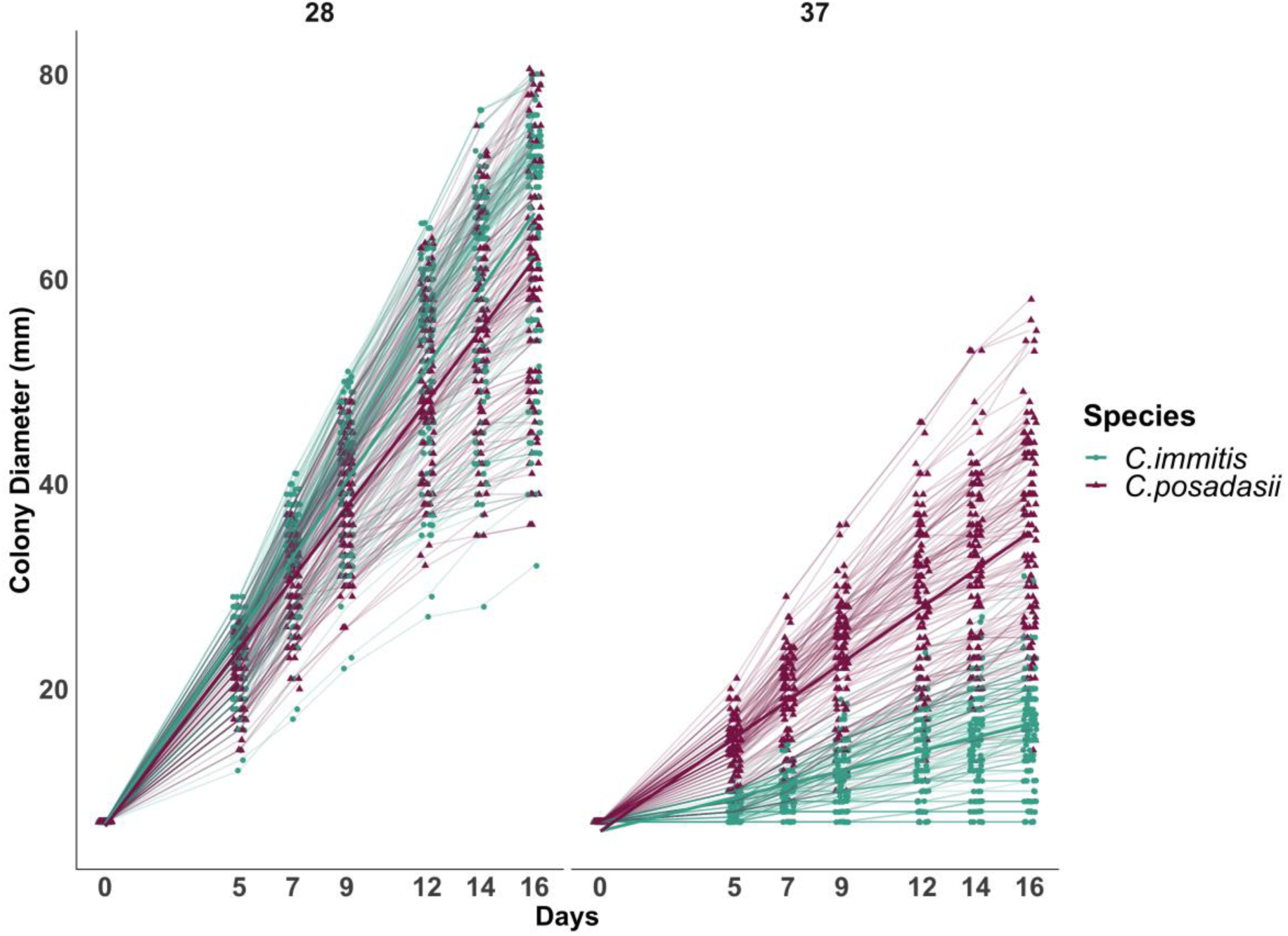
Radial growth rate of 85 isolates of *Coccidioides* demonstrates species-specific response to temperature. Each line represents the mean diameter (y-axis) for each isolate in triplicate (46 *C. immitis* and 39 *C. posadasii*) at a given time point (x-axis). Dark lines represent mean growth rate of each species. Radial growth was measured at day 5, 7, 9, 12, 14 and 16. There is a significant difference in growth rate (slope) in response to higher temperature between species of *Coccidioides*. The radial growth rate of *C. immitis* is decreased at a higher temperature 37°C (slope37 = 0.64 mm/day; 95% C.I. 0.51-0.78) compared to *C. posadasii* (slope37 = 1.82 mm/day; 95% C.I. 1.49-2.16). Both species appear to tolerate 28°C and grow at a similar rate (C. *immitis* slope28 = 3.73 mm/day; 95% C.I. 3.53-3.92, *C. posadasii*, slope28 = 3.47 mm/day; 95% C.I. 2.98-3.90).

### 3.1 Figures, Tables and Schemes

Fig 1. Temperature impacts growth ability of *C. immitis* isolates compared to *C. posadasii* on YE media. Seven mm diameter plugs were sub-cultured onto yeast extract plates and radial growth was documented over 16 days. (A) Radial growth measurements at 37°C for 46 *C. posadasii* and 39 *C. immitis* isolates in triplicate. (B) Radial growth measurements at 28°C for 46 *C. posadasii* and 39 *C. immitis* isolates in triplicate. (C) Representative samples of phenotypic variation observed between species on day 16.

Fig 2. Radial growth rate of 85 isolates of *Coccidioides* demonstrates species-specific response to temperature. Each line represents the mean diameter (y-axis) for each isolate in triplicate (46 *C. immitis* and 39 *C. posadasii*) at a given time point (x-axis). Dark lines represent mean growth rate of each species. Radial growth was measured at day 5, 7, 9, 12, 14 and 16. There is a significant difference in growth rate (slope) in response to higher temperature between species of *Coccidioides*. The radial growth rate of *C. immitis* is decreased at a higher temperature 37°C (slope37 = 0.64 mm/day; 95% C.I. 0.51-0.78) compared to *C. posadasii* (slope37 = 1.82 mm/day; 95% C.I. 1.49-2.16). Both species appear to tolerate 28°C and grow at a similar rate (C. *immitis* slope28 = 3.73 mm/day; 95% C.I. 3.53-3.92, *C. posadasii*, slope28 = 3.47 mm/day; 95% C.I. 2.98-3.90).

## 4. Discussion

Although many studies have looked at genetic variation among isolates of both species of *Coccidioides*, few studies have compared phenotypic differences. Observed genetic diversity between and within species makes it reasonable to hypothesize that phenotypic variation exists. We propose that a methodical documentation of phenotypic variation is a necessary first step to determine the ecological or clinical relevance of these traits. In this study, we have identified a definitive phenotypic difference with a congruent analysis at two institutions for a diverse set of isolates. A total of 85 isolates covering the geographic range of both species show that *C. posadasii* isolates grow at a significantly faster rate (p<0.001, Fig 2 and Table 2) than *C. immitis* isolates in the mycelial form at 37°C on YE agar. Additionally, *C. immitis* grows slightly faster than *C. posadasii* at 28°C on YE agar although the difference in growth rate is not significant (p-value = 0.072, Fig 2 and Table 2). We note that growth rate may be influenced by nutrition source, and the results are limited to the media utilized for the current study.

Functionally, this phenotype is similar to a classic temperature sensitive (ts) conditional mutant, such that *C. immitis* exhibits normal growth at permissive temperature, and significantly slower growth under stressful conditions. It is possible that *C. immitis* could be restored to normal growth at 37°C by gene replacement with appropriate *C. posadasii* alleles if candidate genes were identified. Several genes and pathways have been described in *Aspergillus fumigatus* related to thermotolerance (54). For example, the observed phenotype could be due to mutations in a heat shock protein (Hsp). Hsps are activated in response to changes in temperature and regulate cellular processes associated with morphogenesis, antifungal resistance, and virulence by triggering a wide array of cellular signaling pathways (53, 67). Hsps are activated by a heat shock transcription factor (Hsf) that acts as a thermosensor, regulating the Hsps at specific growth temperatures (68). Several studies have shown that *Coccidioides* up-regulates heat shock proteins Hsp20 and Hsp9/12 at high temperature during the parasitic lifecycle while down-regulating Hsp30 and Hsp90 (69–72). Further investigation of Hsps and Hsfs in *Coccidioides* could elucidate mechanisms of the species-specific thermotolerant behavior observed in this study. Alternatively, many classical ts mutants occur in genes required for normal cellular growth and are due to single amino acid changes that affect protein function or stability at the restrictive temperature. For example, a number of colonial temperature sensitive (*cot*) mutants have been identified in *Neurospora crassa.* The *N. crassa cot-1* mutant has been studied in greatest detail, and the ts defect is due to a SNP causing a single amino acid change in a Ser/Thr protein kinase required for normal hyphal extension, thus resulting in restricted growth at normally permissive temperatures above 32°C (73, 74). Finally, recent work in *Saccharomyces* indicates that mitochondrial genotypes are associated with heat tolerance (75). The mitochondrial genomes of the two species of *Coccidioides* are also distinct, and thus mitochondrial function is another potential mechanism controlling thermotolerance in *Coccidioides*.

The source of the genotypic variation driving the observed phenotype may be attributable to a stochastic event, such as a founder effect or population bottleneck 10-12 MYA, which is the estimated time the two species have been separated (6, 76). Alternatively, the observed pattern may be due to selection pressure from a specific environment, host, or directly associated with virulence. Thus, the observed differential thermotolerance may relate to the saprobic phase of the lifecycle and reflect adaptation to specific environments. A pattern of alternating wet-dry conditions has been related to Valley fever incidence across the southwestern U.S. (5, 77–81). It has been proposed that fungal growth occurs during brief periods of heavy moisture during monsoon and winter rainy seasons in the Southwest, which are followed by prolific conidia production when warm temperatures and low rainfall desiccate soils and increase dispersal via dust (the “grow and blow” hypothesis) (27, 78, 82). Additionally, during high temperature periods, it is hypothesized that the surface soil is partially sterilized and many competitors are removed, but Coccidioides spores remain viable (26). Another hypothesis is that C. posadasii may be better adapted to growth in the high soil temperatures observed in the southwestern deserts compared to the California endemic C. immitis. Maricopa, Pinal and Pima counties harbor the highest coccidioidomycosis case rates in Arizona due to C. posadasii, and according to the National Centers for Environmental Information (83), the annual mean temperature (1901–2000) were 20.7°C, 19.8°C and 19.2°C, respectively. On the other hand, Fresno, King and Kern counties, which harbor the highest coccidioidomycosis case rates in California due to C. immitis, had annual mean temperatures of 12.4°C, 16.9°C and 15.8°C, respectively. The difference in 100-year average annual mean temperature between highly endemic areas of Arizona and California supports our hypothesis that C. posadasii is more adapted to higher temperatures compared to C. immitis. Alternatively, a preferred host species may vary in normal body temperature, in accordance with the endozoan small mammal reservoir hypothesis proposed by Barker and Taylor (84). Interestingly, a decline in mean human body temperature (∼1.6%) has recently been reported (85). Whether this impacts coccidioidomycosis rates is unknown.

Published literature to date suggests that disease outcomes are related primarily to host-specific factors (37, 38, 86), and certainly, host genetic background can impact disease progression. We propose that pathogen-specific variation may also contribute to capricious disease outcomes in coccidioidomycosis patients. Currently, species-specific virulence is not well-documented in *Coccidioides* research, but has been suggested (1, 87). This is in part due to the use of a few characterized laboratory strains of *Coccidioides* for most hypothesis testing, primarily strains Silveira, C735 and RS (70, 86, 88–91). Therefore, connecting phenotypic dissimilarity to established genetic variation using genome-wide association studies could provide insight into unique characteristics of these genetically distinct pathogens.

## 5. Conclusions

In summary, we have identified a significant phenotypic difference between *C. immitis* and *C. posadasii.* Although growth rate on YE media at two temperatures is the only characteristic we explicitly tested, there are certain to be more phenotypic differences between species, and possibly between populations. This, coupled with the recent availability of the genome sequence of multiple strains for both fungal species, may allow comparative genomic approaches to elucidate candidate genes for thermotolerance regulation in *Coccidioides* and closely related Onygenales (7).

## Author Contributions

H.L.M., P.L.H., M.M.T., C.S.W., M.J.O. and B.M.B. prepared the initial draft of the manuscript. B.M.B., M.J.O., and J.N.G. developed the concept, provided funding, and were responsible for approving the final draft of the manuscript. M.M.T., H.L.M., P.L.H. assisted with creation of figures and writing final manuscript. I.N.S, H.L.M., and C.S.W. developed statistical models. H.L.M., P.L.H., B.M.B., performed experiments and collected data. N.P.W, G.R.T. III., R.M-S., L.R.C-O, P.K., C.P., J.T., J.N.G., provided isolates. All authors have read and agreed to the published version of the manuscript.

## Supplementary Materials

Fig S1. Growth of *C. immitis* and *C. posadasii* on YE media at NAU and UA. Seven mm diameter plugs were sub­cultured onto yeast extract plates and radial growth was documented over sixteen days. (A) Radial growth measurements at 28°C and 37°C for 85 isolates in triplicate, at both institutions. (B) Representative samples of phenotypic variation observed between species on day sixteen for both NAU and UA experiments.

Table S1. Analysis of variance. Impact of institution collection site was investigated for each temperature specific model. The factors “day”, “species”, and “lab” were assumed to be fixed linear effects, and individual isolate response for each day was considered to be a normally distributed random effect as appropriate in a longitudinal study. The Factor “Lab” location adds variation to the data sets but does not alter overall findings. Species specific behavior based on temperature pattern remain.

Table S2. Mean Colony Diameter at 28^e^C. Mean diameter and standard deviation for *C. posadasii* and *C. immitis* at 28°C. Welch’s t-test was used to compare difference in means. Mean is significantly different on all days tested. Significance is reduced on day 16.

Table S3. Mean Colony Diameter at 37°C. Mean diameter and standard deviation for *C. posadasii* and *C. immitis* at 37°C. Welch’s t-test was used to compare difference in means at each time point. Mean is significantly different on all days tested.

Table S4. Table. Comparison Linear Model and Bootstrap Values. Comparison of linear model and bootstrap 95% confidence intervals for 28°C and 37°C data sets. Bootstrapping conducted using the boot package in R. S1 File. Final Raw Data for Temperature Differences at 37 °C and 28 °C. Measurements (diameter in mm) for each isolate on each plate were recorded on days 5, 7, 9, 12, 14, and 16. Three replicates were completed for each strain for both temperature conditions. Strain details are listed in Table 1.

## Acknowledgments

We are indebted to G. Koenig at Roche Molecular Systems for her assistance with obtaining several strains for phenotypic analysis. Special appreciation goes to E. Kellner, H. Paes and E. Temporini for their helpful suggestions.

---

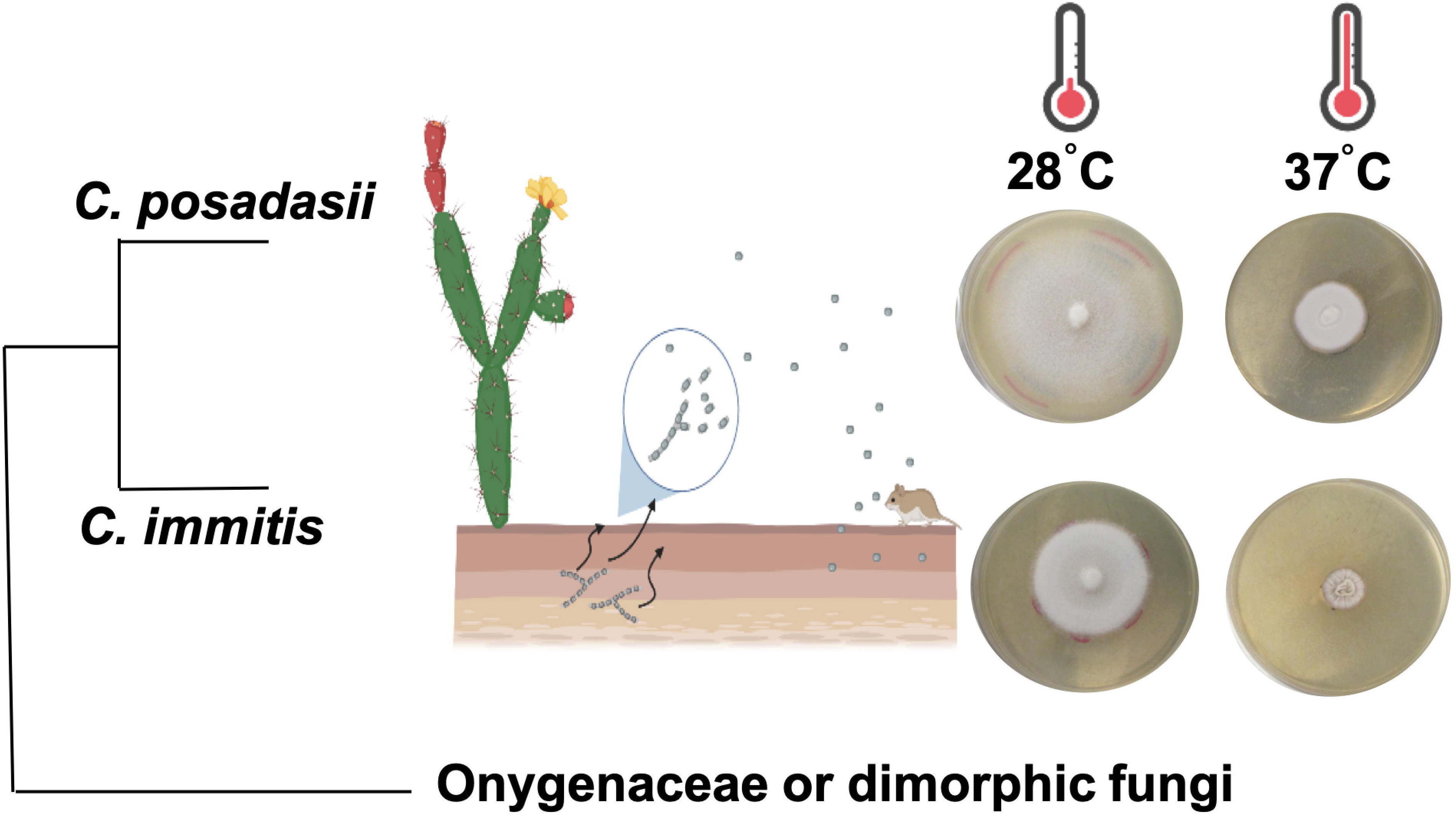

