## supplemental for "Differential thermotolerance adaptation between species of *Coccidioides*": Table S2.docx

Table S2. Mean Colony Diameter at 28ºC

|  | *C. immitis* | | | | *C. posadasii* | | |
| --- | --- | --- | --- | --- | --- | --- | --- |
| *Day* | *Mean diameter (mm)* | *SD* | |  | *Mean diameter (mm)* | *SD* | *p^a^* |
| *5* | 23.66071 | 3.417195 |  | | 22.02564 | 3.308066 | **<0.001** |
| *7* | 33.05357 | 4.462754 | |  | 30.74786 | 4.437606 | **<0.001** |
| *9* | 41.70000 | 5.961447 | |  | 38.91880 | 5.701617 | **<0.001** |
| *12* | 52.81429 | 8.591518 | |  | 49.37179 | 7.860901 | **<0.001** |
| *14* | 59.49643 | 10.140824 | |  | 55.52564 | 9.516512 | **<0.001** |
| *16* | 64.85357 | 11.340622 | |  | 60.63248 | 11.194864 | **<0.01** |
| ^a^Welch’s t-test | | | | |  | | |

Mean diameter and standard deviation for *C. posadasii* and *C. immitis* at 28ºC. Welch’s t-test was used to compare difference in means. Mean is significantly different on all days tested. Significance is reduced on day 16.
