## supplemental for "Differential thermotolerance adaptation between species of *Coccidioides*": Table S3.docx

Table S3. Colony Diameter at 37ºC

|  | *C. immitis* | | | | *C. posadasii* | | |
| --- | --- | --- | --- | --- | --- | --- | --- |
| *Day* | *Mean diameter (mm)* | *SD* | |  | *Mean diameter (mm)* | *SD* | *p^a^* |
| *5* | 8.509929 | 1.098525 |  | | 13.717949 | 3.020322 | **<0.001** |
| *7* | 10.156028 | 1.730127 | |  | 18.512821 | 4.656966 | **<0.001** |
| *9* | 11.812057 | 2.706170 | |  | 22.824786 | 5.814413 | **<0.001** |
| *12* | 14.088652 | 3.798629 | |  | 28.632479 | 7.547377 | **<0.001** |
| *14* | 15.663121 | 4.468739 | |  | 31.882906 | 8.470428 | **<0.001** |
| *16* | 16.734043 | 5.129242 | |  | 35.068376 | 9.421158 | **<0.001** |
| ^a^Welch’s t-test | | | | |  | | |

Mean diameter and standard deviation for *C. posadasii* and *C. immitis* at 37ºC. Welch’s t-test was used to compare difference in means at each time point. Mean is significantly different on all days tested.
