## supplemental for "Differential thermotolerance adaptation between species of *Coccidioides*": Table S4.docx

**S4 Table. Comparison of Linear Model and Bootstrap Values.**

|  | **Linear Model Estimates** | | **Bootstrap Estimates** | |
| --- | --- | --- | --- | --- |
| *Species* | *Mm/days* | *CI 2.5% 97.5%* | | *CI 2.5% 97.5%* |
| *C. immitis* x Day 28ºC | 3.73 | 3.53 – 3.92 | | 3.547 – 3.922 |
| *C. posadasii x* Day 28ºC | 3.47 | 2.98 – 3.9 | | 2.961 – 3.909 |
| *C. immitis* x Day 37ºC | 0.64 | 0.51 – 0.78 | | 0.510 – 0.780 |
| *C. posadasii x* Day 37ºC | 1.82 | 1.49 – 2.16 | | 1.5 – 2.161 |
| N^a^ | 85 | S^b^ | | 2000 |
| ^a^ number of isolates | ^b^ number of simulations | |  | |

Comparison of linear model and bootstrap 95% confidence intervals for 28ºC and 37ºC data sets. Bootstrapping conducted using the boot package in R.
