## Supplementary figures and images for "Differential thermotolerance adaptation between species of *Coccidioides*"

### S1Fig.tiff

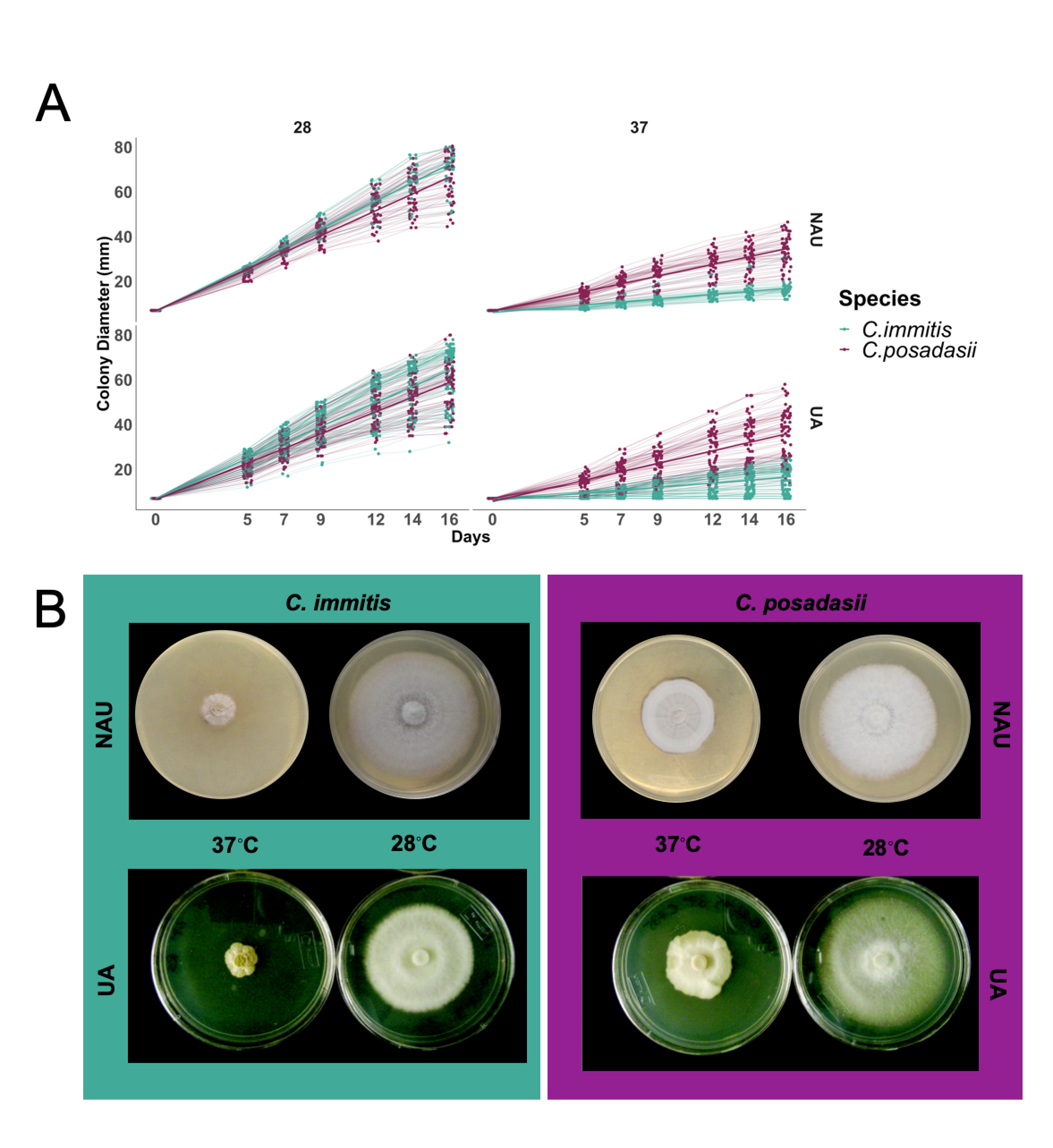
